# Detection of alternative splicing in *Diabrotica virgifera virgifera* LeConte, in association with Bt resistance using RNA-seq and PacBio Iso-Seq

**DOI:** 10.1101/2020.08.03.234682

**Authors:** Zixiao Zhao, Christine G. Elsik, Bruce E. Hibbard, Kent S. Shelby

**Affiliations:** Division of Plant Sciences, University of Missouri, Columbia, MO, USA; Division of Animal Sciences, University of Missouri, Columbia, MO, USA; Institute for Data Science and Informatics, University of Missouri, Columbia, MO, USA; USDA-ARS Plant Genetics Research Unit, Columbia, MO, USA; USDA-ARS Biological Control of Insects Research Laboratory, Columbia, MO, USA

**Author notes:** Author contact information ZZ; CGE; BEH; KSS.

**Keywords:** western corn rootworm, alternative splicing, Bt resistance, Iso-seq

## Abstract

**Background:** Alternative splicing is one of the major mechanisms that increases transcriptome diversity in eukaryotes, including insect species that have gained resistance to pesticides and Bt toxins. In western corn rootworm (*Diabrotica virgifera virgifera* LeConte), neither alternative splicing nor its role in resistance to Bt toxins has been studied.

**Results:** To investigate the mechanisms of Bt resistance we carried out single-molecule real-time (SMRT) transcript sequencing and Iso-seq analysis on resistant, eCry3.1Ab-selected and susceptible, unselected, western corn rootworm neonate midguts which fed on seedling maize with and without eCry3.1Ab for 12 and 24 hours. We present transcriptome-wide alternative splicing patterns of western corn rootworm midgut in response to feeding on eCry3.1Ab-expressing corn using a comprehensive approach that combines both RNA-seq and SMRT transcript sequencing techniques. We found that 67.73% of multi-exon genes are alternatively spliced, which is consistent with the high transposable element content of the genome. One of the alternative splicing events we identified was a novel peritrophic matrix protein with two alternative splicing isoforms. Analysis of differential exon usage between resistant and susceptible colonies showed that in eCry3.1Ab-resistant western corn rootworm, expression of one isoform was significantly higher than in the susceptible colony, while no significant differences between colonies were observed with the other isoform.

**Conclusion:** Our results provide the first survey of alternative splicing in western corn rootworm and suggest that the observed alternatively spliced isoforms of peritrophic matrix protein may be associated with eCry3.1Ab resistance in western corn rootworm.

## Introduction

The western corn rootworm (WCR), *Diabrotica virgifera virgifera* LeConte (Coleoptera: Chrysomelidae) is one of the most destructive maize pests in the US Corn Belt. WCR has demonstrated the genetic plasticity to develop resistance to many control strategies, including transgenic maize expressing rootworm-targeting Bt toxins Cry3Bb1, mCry3A, eCry3.1Ab, and Cry34/35Ab1. Resistance to Bt maize has been selected in laboratory colonies (Frank et al. 2013; Meihls et al. 2011; Meihls et al. 2008). In the field, failure of Bt-maize lines to control rootworm has been reported in multiple states (Gassmann et al. 2011; Ludwick et al. 2017; Wangila et al. 2015; Zukoff et al. 2016). This has highlighted the importance of understanding the mechanisms of WCR Bt resistance to better inform management strategies.

The mechanisms of Bt resistance in WCR are not well elucidated at the gene level. High-throughput RNA sequencing (RNA-seq) provides a powerful tool to analyze gene expression profiles, especially to identify differentially expressed genes that are associated with Bt resistance. Transcript sequences generated in previous studies were either from expressed sequence tags (EST) or assembled *de novo* from short-length RNA-seq reads (Rault et al. 2018; Wang et al. 2017; Zhao et al. 2019). In other studies, genetic markers were developed for analyzing quantitative traits loci (QTL) that were associated with Bt resistance in WCR (Flagel et al. 2015). However, gene expression, regulation and genomic variation may not be the only molecular phenomena associated with resistance. Post-transcriptional processes such as transcript alternative splicing (AS) may result in isoforms that could confer additional functions, *i.e.* Bt resistance, in WCR.

AS is a common phenomenon in eukaryotes. A multi-exon gene can be processed utilizing alternative splice sites to create multiple resulting transcripts with different exon combinations. In humans, it has been estimated that about 95% of multi-exon genes produce more than one transcript by AS (Pan et al. 2008). In insects, AS is involved in many physiological processes, including sex determination (Salz 2011), social behaviors (Jarosch et al. 2011), molting (Abdel-Banat and Koga 2002) and immune responses (Riddell et al. 2014). AS of some insecticide receptors such as sodium channels can produce receptor proteins with decreased binding site sensitivity, resulting in insecticide resistance (Dong 2007). Glutathione transferases, a class of detoxification enzyme involved in resistance to insecticides, are alternatively spliced in some mosquito species (Kasai et al. 2009; Strode et al. 2008). Recent studies on the pink bollworm (*Pectinophora gossypiella*) demonstrated that splice variants of a cadherin receptor *PgCad1* and an ATP-binding cassette transporter (ABC transporter) *Pg*ABCA2 caused resistance to Cry1Ac and Cry2Ab (Fabrick et al. 2014; Mathew et al. 2018). Similarly, AS of corn earworm (*Helicoverpa armigera*) ABC transporter *Ha*ABCC2 was shown to be associated with Cry1Ac resistance (Xiao et al. 2014). These studies suggest that AS could produce variants allowing rapid adaptation to biotic stressors such as Bt intoxication.

No previous studies have focused on the role of AS in WCR. One limitation is that the WCR genome is complicated and less annotated compared to other studied species. The *de novo* assembled transcripts from short read RNA-seq do not provide an accurate representation of transcript isoforms. To circumvent this difficulty we have adopted the newly introduced long read single molecule real-time (SMRT) technology of Pacific Biosciences (PacBio) (Rhoads and Au 2015). This technology generates high-quality, ultra-long sequence reads greater than 10 kb. Isoform sequencing (Iso-seq) is the application of PacBio sequencing to transcriptomes, which can produce full-length transcripts and splicing variants without assembly artifacts such as chimeras (Gonzalez-Garay 2016). PacBio Iso-seq provides simplified and reliable approaches for detecting AS that can improve WCR genome annotation.

We report here the first genome-wide survey of splicing isoforms in WCR using PacBio Iso-seq. The isoform profile was generated from a lab-selected eCry3.1Ab-resistant colony and its non-selected susceptible counterpart colony (Frank et al. 2013). To detect the AS events associated with Bt resistance, we conducted a comprehensive analysis from a previous Illumina-based RNA-seq study of midgut response to eCry3.1Ab (Zhao et al., 2019), with the longer transcripts provided by PacBio Iso-seq. We were able to identify differentially spliced transcripts between eCry3.1Ab-resistant and susceptible WCR colonies, with or without intoxication from the eCry3.1Ab. Our results revealed that for many genes, resistant and susceptible WCR have different splicing patterns. Most of those patterns are specific to Bt intoxication and exposure time. However, a few genes displayed colony-specific patterns regardless of intoxication and exposure time. The results herein will provide valuable resources for understanding the genetic and molecular basis of Bt resistance in WCR.

## Results

### WCR transcriptome Iso-Seq and bioinformatics pipeline

In order to increase the sequence coverage, we pooled high-quality total RNA extracted from eight WCR midgut samples. The larvae of WCR were recovered from bioassays where the neonates from eCry3.1Ab-resistant and -susceptible colonies were fed with the seedling roots of eCry3.1Ab-expressing and non-Bt isoline maize lines for 12 and 24 hours (Table S1). A SMRT library was prepared from pooled total RNA from all treatments and sequenced in three SMRT cells. A total of 1,046,323 circular consensus sequences (CCS) were generated, from which 745,428 full-length non-chimeric sequences were generated in the classification step. After the clustering and polishing step, 53,394 high-quality full-length sequences and 322,095 low-quality full-length sequences were generated. The alignment of both high-quality and low-quality full-length sequences to the *D. virgifera virgifera* genome assembly v2 showed that 37,944 high quality (71.06%) and 195,768 low quality (60.78%) sequences could be aligned.

### WCR gene loci construction and isoform detection

A bioinformatics pipeline was developed using publicly available tools (Figure 1). We first merged the high-quality and low-quality full-length sequences that aligned to the *D. virgifera virgifera* genome. Then the post-analysis script Cupcake ToFU (https://github.com/Magdoll/cDNA_Cupcake/wiki/Cupcake-ToFU:-supporting-scripts-for-Iso-Seq-after-clustering-step) was used to collapse transcripts, resulting in a total of 11,662 gene loci and 34,081 isoforms. Among those there were 3,851 single-exon genes. For 7,811 multi-exon gene, 5,291 (67.73%) of them produced more than one transcript.

**Figure 1.**
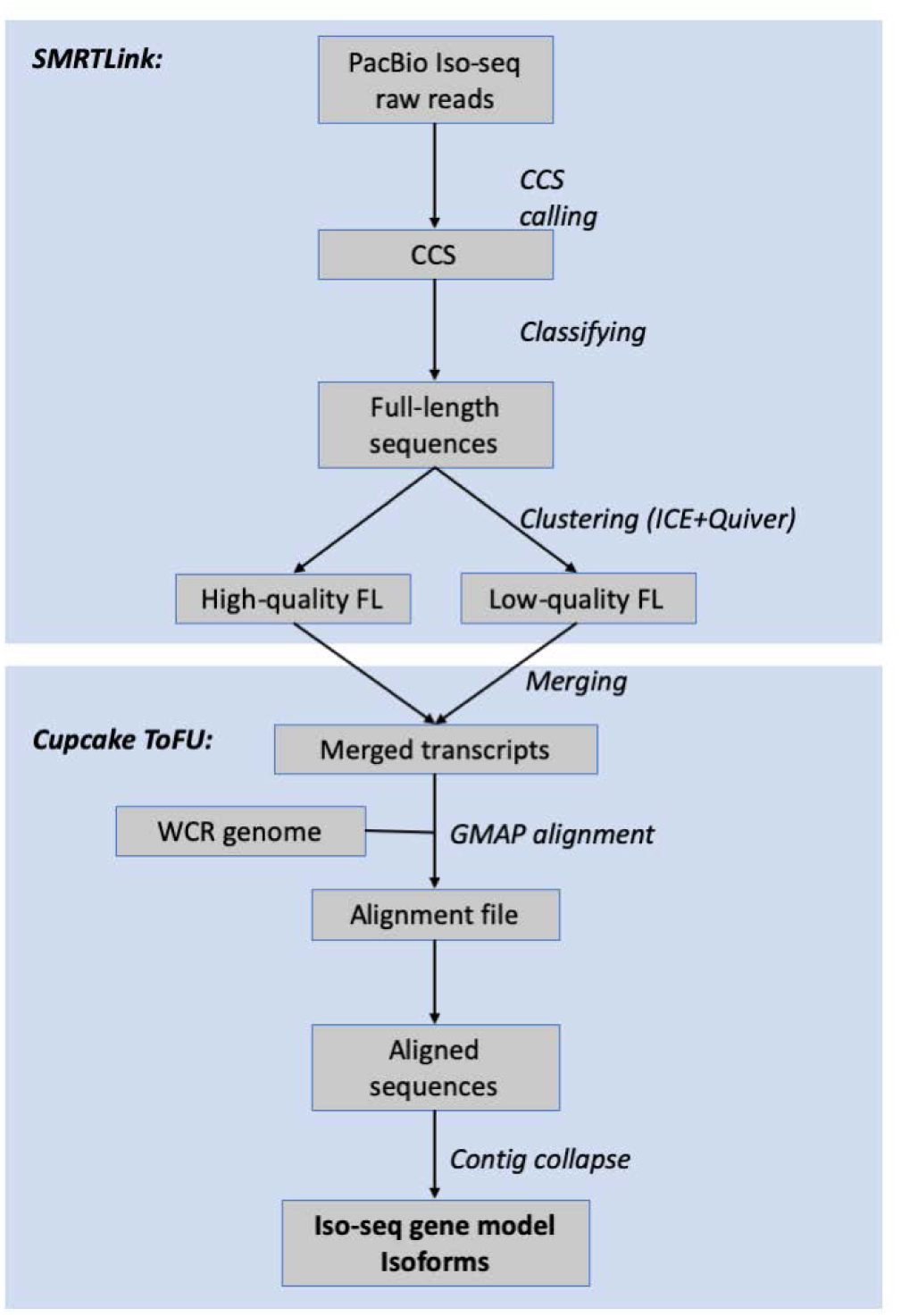
Bioinformatics pipeline of PacBio Iso-seq data analysis and the construction of gene models.

BUSCO (Simão et al. 2015) was adopted to evaluate the completeness and integrity of the Iso-seq isoforms. From the insect conserved single copy orthologs dataset provided by OrthroDB, 63.7% of the isoforms were identified as complete transcripts. There were 6.7% incomplete and 29.6% missing orthologs (Table 1), which indicated that some ortholog genes in OrthroDB were not represented in this Iso-seq isoform dataset.

**Table 1.** Statistics and BUSCO analysis (version v3) results of WCR midgut isoforms from PacBio Iso-seq.

|  |  |
| --- | --- |
| **Total isoform contig number** | 34,081 |
| **N50** | 2,072 |
| **Complete BUSCO orthologs** | 63.70% |
| Orthologs reported as single-copy in isoforms | 41.40% |
| Orthologs reported as duplicated in isoforms | 22.30% |
| **Fragmented BUSCO orthologs** | 6.70% |
| **Missing BUSCO orthologs** | 29.60% |

We compared the gene models created based on Iso-seq alignment with RefSeq automated annotation using GFFcompare (Pertea and Pertea 2020). Iso-seq revealed a higher average AS rate (2.5 isoforms per locus) than RefSeq prediction (1.3 isoforms per locus). There were 7,117 loci which appeared in both Iso-seq and RefSeq models, while 5,081 and 17,885 loci appeared only in Iso-seq or RefSeq models, respectively. We also found some Iso-seq loci that were split in RefSeq, suggesting that Iso-seq can be used to correct some of the RefSeq models.

### Functional annotation of WCR Iso-seq transcriptome

All isoforms and transcripts were functionally annotated by Blast2GO (Conesa et al. 2005) using the standard pipeline. BLASTX against the NCBI non-redundant protein database (nr) showed 28,801 (84.5%) of Iso-seq isoforms had significant hits (E-value < 1e-3). Of these, 20,027 (57.8%) could be annotated with GO terms. The direct counts of GO terms of annotated isoforms showed that the most abundant GO terms of biological process (BP), molecular function (MF) and cellular component (CC) were oxidation-reduction process (GO:0055114), ATP binding (GO:0005524) and integral component of membrane (GO:0016021). The distribution of GO terms was correlated with the primarily digestive function of insect midgut (Figure S2).

As a result of functional annotation, the most highly spliced gene locus in Iso-seq model (PB.6253) was predicted as a mediator of RNA polymerase II transcription subunit 12 (MED12) with 170 AS isoforms. We were able to identify the splicing pattern of potential Bt receptors cadherin and aminopeptidase N (APN). The cadherin (EF531715), previously identified in WCR (Sayed et al. 2007), had 17 different AS isoforms (PB.2010). There was another putative DE-cadherin (PB.6835) gene with 9 isoforms, and one putative cadherin-99C (PB.5905) with one isoform from the WCR genome. We also identified 5 different APN genes (PB.8995, PB.9388, PB.721, PB.797, and PB.6333), each with 2 to 13 isoforms (Table 2).

**Table 2.** Alternative splicing patterns of cadherin-like proteins and Aminopeptidase N.

| Iso-seq gene ID | Description | Genome Contig | Number of isoforms |
| --- | --- | --- | --- |
| PB.2020 | Cadherin-like protein | NW_02103863 | 18 |
| PB.6835 | DE-cadherin | NW_021039636 | 9 |
| PB.5905 | Cadherin-99C | NW_021039636 | 1 |
| PB.8995 | Aminopeptidase N-like protein | NW_021040963 | 5 |
| PB.9388 | Aminopeptidase N | NW_021041366 | 7 |
| PB.721 | Aminopeptidase N-like protein | NW_021038542 | 13 |
| PB.797 | Aminopeptidase N-like protein | NW_021038547 | 2 |
| PB.6333 | Aminopeptidase N-like protein | NW_021039468 | 5 |

### Detection of the differential exon usage between resistant and susceptible colonies

To analyze whether AS is associated with eCry3.1Ab resistance, we developed a pipeline that leveraged Illumina RNA-seq data from the same colonies and treatments (Zhao et al. 2019) to determine differential exon usage (DEU). The DEXSeq (Reyes et al. 2012) package was adopted to report gene loci from Iso-seq with DEU between resistant and susceptible colonies under different feeding treatments. For colonies that fed on eCry3.1Ab-seedlings, DEXSeq reported 35 loci with DEU at 12 hours, and 60 loci with DEU at 24 hours. For colonies that fed on isoline seedlings, DEXSeq identified 47 loci with DEU at 12 hours and 90 loci with DEU at 24 hours. The relationship of those loci is shown in a Venn diagram (Figure 2). Four loci (PB.9337, PB.11067, PB.8408, and PB.6428) had DEU exons between colonies in all treatments. Each of them had isoforms ranging from 2 to an extreme of 23. Functional annotation showed the predicted functions of those loci were actin-related protein 2/3 complex subunit 4 (PB. 9337), mitochondrial ATP synthase subunit gamma (PB.111067), 60S ribosomal protein L10a (PB.8408) and peritrophic membrane protein (PMP) containing chitin-binding domain cbd-1 (PB.6428) (Table 3). To characterize the functions of those loci with DEU, we applied GO enrichment analysis using Fisher’s exact test. The results demonstrated that under different experimental conditions, genes with DEU had unique enriched GO terms. None of the GO terms were significantly enriched for all four experimental conditions, as shown in the Venn diagram (Figure 3). However, when being fed with eCry3.1Ab-maize root, some GO terms related to detoxification (GO:0098754, GO:0061687, and GO:1990461), ion transportation (GO:0098659, GO:0097286, GO:0098711, GO:0097577, GO:0087459, and GO0099587), and response to superoxide (GO:0019430, GO:0071450, and GO:0000303) were enriched at both 12 and 24 hours after feeding, which demonstrated the functions of AS genes in response to eCry3.1Ab intoxication (Table S3).

**Table 3.** Gene loci with differential exon usage between colonies in all experimental conditions.

| Gene Loci | Isoforms | Exon (flattened) involved under experimental conditions |  |  |  |
| --- | --- | --- | --- | --- | --- |
|  |  | Bt fed, 12 h | Bt fed, 24 h | Isoline fed, 12 h | Isoline fed, 24 h |
| PB.6428 | 2 | E001/E002 | E001/E002 | E001/E002/E003 | E001/E002/E003 |
| PB.8408 | 9 | E001/E008/E013 | E001/E013 | E013 | E010/E011/E013 |
| PB.9337 | 7 | E012 | E003/E004 | E004/E002 | E003/E004/E008 |
| PB.11067 | 23 | E030/E032 | E032 | E031/E032 | E030/E031 |

**Figure 2.**
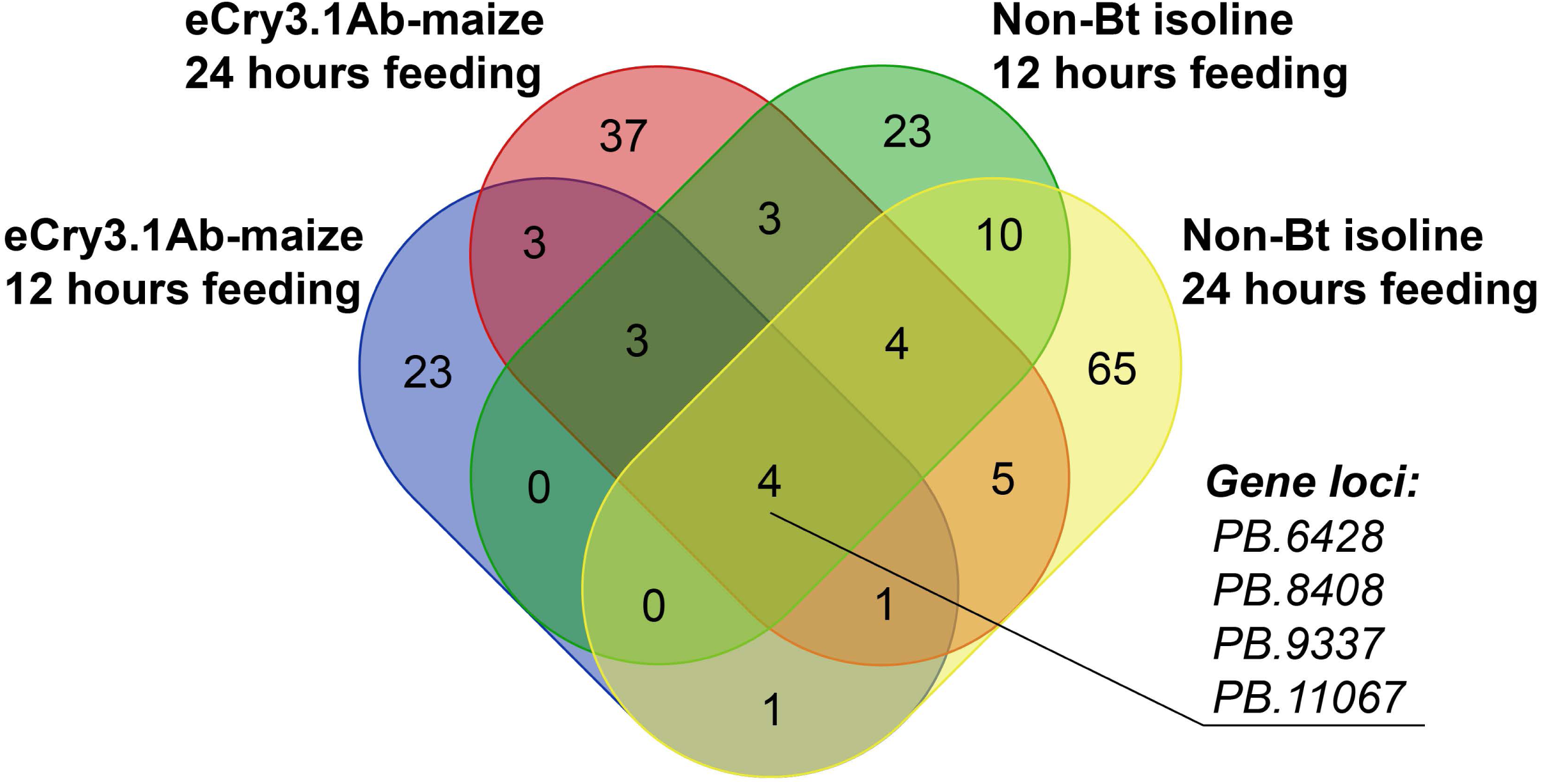
Venn diagram of WCR midgut gene loci with differential exon usage (DEU) between eCry3.1Ab-resistant and susceptible colonies after feeding with eCry3.1Ab-expressing or non-Bt isolines maize respectively for 12 and 24 hours. Four gene loci: PB.6428, PB.8408, PB.9337 and PB.11067 displayed DEU at all four experimental conditions.

**Figure 3.**
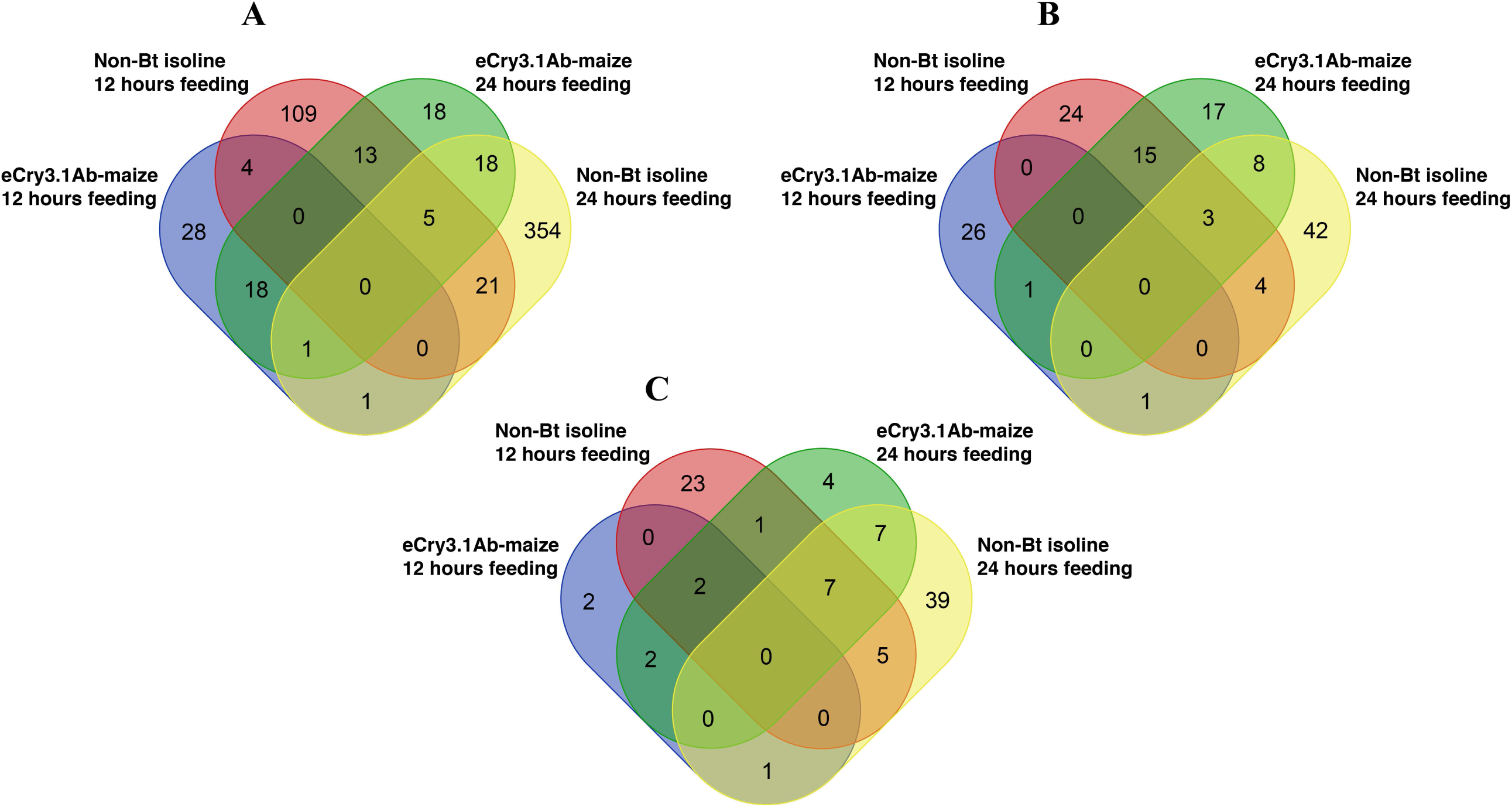
Venn diagram of enriched GO terms of WCR midgut genes with DEU after feeding with eCry3.1Ab-expressing and non-Bt isoline maize respectively for 12 and 24 hours. None of GO terms were enriched in all four experimental conditions. GO term category: **A** biological process (BP); **B** molecular function (MF); **C** cellular compartment (CC).

### Characterization of WCR genes with DEU between resistant and susceptible colonies

Molecular interactions between Coleoptera-targeting Cry3 toxins and midgut cells and resistance mechanisms are still poorly understood in WCR. The differences in splicing patterns between eCry3.1Ab-resistant and -susceptible colonies may be the result of selection pressure imposed by eCry3.1Ab intoxication over many generations. The gene PB.6428, with a predicted function of a novel PMP, contains four exons and two isoforms with alternative promoters of either exon 1 or exon 3 (Figure 4). The pattern was confirmed by PCR using primers flanking the splicing events (see Figure S4, Additional file 4). In all four treatments, the usage of exons 1 and 2 was significantly higher in eCry3.1Ab-resistant WCR larval midguts, while the usage of exon 3, if not significantly, was slightly higher in susceptible larval midguts (Figure 4). This result suggests both isoforms existed in both colonies, but that eCry3.1Ab-resistant WCR tended to use isoform PB.6428.2 while susceptible WCR tended to use PB.6428.1. Further analysis showed the sizes of PB.6428 corresponding peptides were either 99 (PB.6428.1) or 112 (PB.6428.2) amino acids with one conserved 6-cysteine ChtBD2-type chitin binding domain (pfam01607) (Figure 5). A 16 amino acid N-terminal signal peptide was predicted from both isoforms (Figure S5), which indicates proteins of both isoforms could be translocated to midgut luminal compartments in order to bind to chitin within the peritrophic membrane.

**Figure 4.**
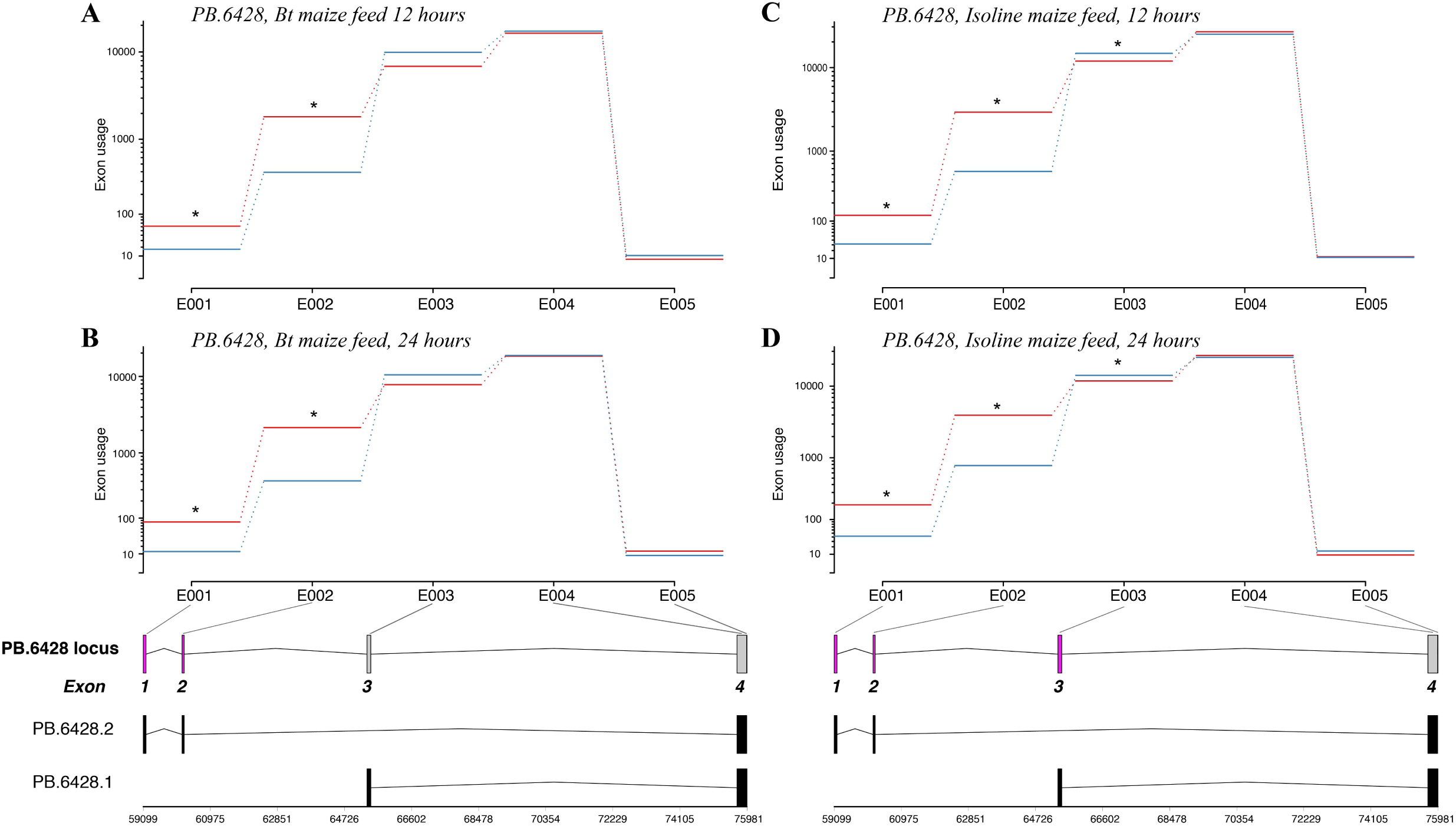
Exon usage of the novel PMP locus PB.6428 in eCry3.1Ab resistant (red) and susceptible (blue) WCR midgut under different feeding conditions: eCry3.1Ab-expressing maize roots for 12 (A) and 24 (B) hours, and non-Bt isoline maize root for 12 (C) and 24 hours (D). The splicing pattern is demonstrated. Exon 4 was flattened and split into E004 and E005 by DEXSeq.

**Figure 5.**
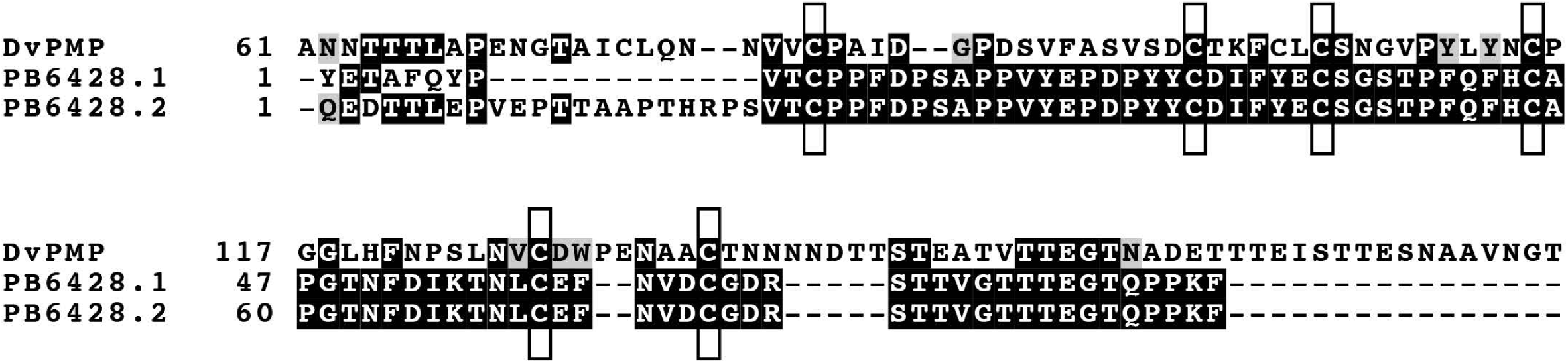
Sequence alignment of predicted protein sequences from PB.6428 isoforms and a predicted WCR peritrophic matrix protein C4 (DvPMP C4, GenBank Accession number: MH256860.1). The conserved 6-cysteine residues of ChtBD2-type chitin binding domain are boxed. The signal peptide of each sequence (see Figure S5, Additional file 5) has been removed.

## Discussion

The mode of action of Bt toxin and the mechanism of Bt resistance has been extensively studied in lepidopteran insects (Bravo et al. 2007; Bravo and Soberón 2008; Heckel et al. 2007). In WCR, the research is restricted by a complicated and incomplete reference genome assembly. The common strategy to overcome this limitation is quantification of genes on a *de novo* assembled transcriptome. The third generation PacBio sequencing technology generates ultra-long reads and serves as a powerful tool to improve genome sequencing, genome annotation, novel gene discovery, and isoform presentation. We implemented the first isoform survey on WCR using PacBio Iso-seq and did a comprehensive analysis in combination with RNA-seq, particularly the splicing profiles in association with eCry3.1Ab resistance. The goal for Iso-seq was to provide an accurate pattern of gene structure of WCR, while RNA-seq data had a higher sequence depth, allowing us to quantify the usage of particular exons in specific colonies and treatments. The genome sequence of WCR allowed us to create gene models based on Iso-seq alignment and to quantify exon usage. However, a large number of Iso-seq transcripts failed to align to the genome, indicating large gaps exist in the current genome assembly.

We carried out differential exon usage analysis on the gene loci identified based on Iso-seq alignment to the reference genome. We discovered that 68% of multi-exon genes in WCR had an AS event, which was significantly greater than for *Drosophila* EST data (19%) (Kim et al. 2006). The result was consistent with the large WCR genome with many transposable elements (TEs) (Coates et al. 2012), which may cause AS events. The DEU exons presented different splicing patterns between samples. When comparing gene loci with DEU between eCry3.1Ab-resistant and susceptible WCR, only a few loci showed consistent DEU regardless of intoxication or feeding time. GO enrichment results showed that for loci with colony-wise DEU, some of their GO terms were enriched, but the enriched terms varied by experimental conditions. The results suggest that AS fine tunes the regulation of gene expression and protein function.

The mis-spliced protein of Bt receptors cadherin and ABC transporters have been found in several resistant Lepidopteran species (Fabrick et al. 2014; Mathew et al. 2018; Xiao et al. 2014). However, the splicing patterns in WCR have not been studied. Using Iso-seq, we documented that some putative Bt receptors, i.e. cadherin (EF531715) (Sayed et al. 2007) and Aminopeptidase N, were alternatively spliced. However, these genes did not show differences in exon usage between eCry3.1Ab-resistant and -susceptible colonies.

In insects, PMP is a group of chitin-binding proteins expressed only in midgut which have one or more chitin binding domains (Tetreau et al. 2015). Although the peritrophic matrix is a barrier to pathogens, Bt toxins can diffuse past this barrier to reach receptors on epithelial cells. The proposed functions for PMP include binding and cross-linking chitin fibrils to form the peritrophic matrix. In Lepidoptera Cry toxins are able to bind PMP, and this binding may contribute to Bt susceptibility in some insects (Rees et al. 2009; Valaitis and Podgwaite 2013). Some potential Bt receptors such as APNs are also located in the peritrophic membrane (Campbell et al. 2008; Hu et al. 2012). In the presented study, we reported a novel PMP locus PB. 6428, which consisted of two AS isoforms. From DEXSeq, the usage of promotor exon 1 and exon 2 of PB.6428.2 were significantly higher in eCry3.1Ab-resistant WCR, regardless of experimental conditions. Meanwhile the usage of promotor exon 3 of PB.6428.1 was slightly lower in in eCry3.1Ab-resistant WCR. These results suggest that eCry3.1Ab-resistant WCR have a bias to express PB. 6428.2 while susceptible WCR to the other isoform. Further studies are necessary to determine whether the alternative splicing of PMP would affect peritrophic matrix stability in WCR midguts, or otherwise show involvement with resistance mechanisms.

In conclusion, we present a survey of alternative splicing patterns in WCR larval midguts using a combination of RNA-seq and PacBio Iso-seq. The full-length assembly-free transcriptome sequences represent a source for annotating gene structures. We also identified gene loci with differential splicing patterns between eCry3.1Ab-resistant and susceptible WCR; one of them is a novel PMP with two isoforms. We briefly characterized this gene, but its functions and specific mechanisms in eCry3.1Ab resistance are yet to be studied.

## Experimental Procedures

### Insects and data sources

The non-diapause eCry3.1Ab-resistant and susceptible WCR colonies were from continuous lab-selection experiments (Frank et al. 2013). After hatching, about 40-50 neonate larvae were transferred to a Petri dish with 3-4 maize seedlings (∼7 days after germination) placed on moistened filter paper. After feeding for 12 or 24 hours (23 °C in dark), surviving larvae were collected for midgut dissection. An RNA-Seq experiment with the same colonies and treatments has been previously described (Zhao et al. 2019) and the sequence data are accessible from the NCBI BioProject database (Accession: PRJNA416935). The *D. virgifera virgifera* reference genome assembly (version 2) and the RefSeq annotation are accessible from the NCBI assembly database (GenBank Assembly Accession: GCA_003013835.2).

### Bioassay, Iso-seq library preparation, and sequencing

Total RNA was extracted from ∼25 dissected neonate midguts using Direct-zol RNA Mini Prep kit (Zymo Research, Irving, CA) with DNase treatment (Thermo Fisher, Waltham, MA). To increase the sequence coverage of Iso-seq we pooled an equal amount of total RNA from each sample. Iso-seq library preparation and PacBio sequencing were performed by Genewiz LLC (South Plainfield, NJ). The library was loaded and sequenced on three cells of PacBio Sequel sequencer to increase the sequence depth. Size-selection was not implemented, according to factory recommendations.

### Iso-seq data processing and transcript analysis

The PacBio SMRTLink v5.0.1 Iso-seq pipeline was employed to extract sequences from sequencer outputs with default settings. The pipeline contained three steps. The first circular consensus sequence (CCS) step generated CCS reads from sequencer outputs. The second classification step identified full-length transcripts. The third clustering step further clustered and polished full-length sequences using ICE and Quiver algorithms and partitioned the full-length sequences into high-quality and low-quality categories. To maximize the sensitivity of detecting gene and isoforms, we merged both high-quality and low-quality full-length transcripts. The post-analysis pipelines were applied following the SMRTLink pipeline to construct gene models and detect isoforms. We first aligned full-length transcripts to the WCR genome assembly using GMAP (version 2014-08-19) (Wu and Watanabe 2005). The aligned transcripts were processed by Cupcake ToFU. A custom Perl script was used to convert the gene models in GFF format to a standard genomic feature file in GTF format. To compare the Iso-seq gene models of aligned transcripts with automated RefSeq annotation, GFFCompare was used with default settings. To evaluate the completeness and coverage of Iso-seq we applied BUSCO (version 3) with the lineage “insect_obd9” on aligned gene isoforms.

### Identification of colony-specific spliced genes

Using RNA-seq reads we compared the differences in alternative splicing patterns between eCry3.1Ab-resistant and susceptible colonies. We first aligned all the RNA-seq reads to the WCR reference genome using TopHat (version 2.0.12) (Trapnell et al. 2009). The DEXSeq package (Anders et al. 2012) was then employed to test changes in exon usage of each gene between resistant and susceptible samples under different experimental conditions. The differential exon usage (DEU) of each gene is reported and represents the relative abundance of each exon within a gene, which demonstrates the differences of isoform patterns between colonies.

### Functional annotation of Iso-seq transcriptome and GO enrichment analysis of gene set with DEU between colonies

We functionally annotated the Iso-seq transcriptome using the Blast2GO (version 5.2.5) standard annotation pipeline. The isoforms were first searched against the NCBI non-redundant protein database (nr) using BLASTX. The gene ontology (GO) terms were obtained by mapping BLASTX results to the Gene Ontology annotation database (GOA) (Huntley et al. 2014). Then we applied Blast2GO with default annotation rules to above GO terms, resulting in the GO annotation with reliability and specificity. We also identified protein domains and motifs of isoforms using InterProScan. The GO terms reported from InterProScan were merged with the Blast2GO GO annotations.

To obtain GO annotations of each gene, we merged all the annotated GO terms from its isoforms. To better understand the function of colony-specific spliced genes we applied GO enrichment analysis using the R package topGO (Alexa and Rahnenfuhrer 2010). The Fisher’s exact tests algorithm was applied separately on three categories of GO terms: biological process (BP), molecular function (MF), and cellular component (CC). The significant GO terms (Fisher’s p-value < 0.05) are reported.

### Characterization of WCR genes with DEU exons between resistant and susceptible colonies

SingleIP-5.0 (Almagro Armenteros et al. 2019) was applied to translated protein sequences to predict the signal peptides. The cDNA of WCR midgut was synthesized by Omniscript RT kit (Qiagen, Germantown, MD). To validate the pattern of AS, PCR primers were designed on flanking regions of splicing sites (see Table S6, Additional file 6) and the products are observed in 1% agarose gel.

## Acknowledgements

This research was funded by the USDA-ARS and the University of Missouri. Maize seeds were provided by Syngenta Biotechnology. We thank Julie Barry, Michelle Gregory, Justin Le Tourneau, and Christopher A. Bottoms for technical and bioinformatic assistance.

